# The visual encoding of familiar and unfamiliar tools

**DOI:** 10.1101/2025.05.24.655951

**Authors:** Luigi Valio, Maria Cuomo, Ciro Rosario Ilardi, Giovanni Federico

## Abstract

How do we extract meaning and understand the action potential of everyday objects? In this eye-tracking study, we show that familiarity with tools modulates the temporal dynamics of their visual exploration. Twenty-five right-handed participants (14 females; mean age = 23.6 ± 3.34 years) freely observed images of familiar and unfamiliar tools while their eye movements were recorded over a 1500-millisecond window. All tools were post-hoc segmented into functional and manipulative areas. Participants initially prioritized functional components—particularly for unfamiliar tools. Attention to manipulative features emerged later and remained attenuated throughout viewing. In contrast, familiar tools prompted earlier and sustained visual attention on manipulative regions, suggesting faster access to action-related representations. These results support a sequential semantic-to-sensorimotor model of object perception, in which previous knowledge facilitates the transition from semantic identification to action-related engagement.

## Introduction

The human capacity to interact with tools reflects the integration of perceptual, semantic, and sensorimotor knowledge (Federico, Osiurak, Reynaud, et al., 2021; Federico, Osiurak, Ciccarelli, et al., 2023; Federico & Brandimonte, 2019, 2020). Over the past fifty years, researchers have extensively debated the role of the action-related features of tools, known as affordances, defined as the perceived opportunities for interaction offered by an object (Gibson, 1979; Osiurak et al., 2017). Central to this debate is the framework asserting that “vision guides action”, which has significantly shaped research on tool-related visual processing (Milner & Goodale, 2008).

Research on tool perception has predominantly focused on sensorimotor aspects, emphasizing the tight coupling between visual perception and motor activation (Bach et al., 2014; Cho & Proctor, 2010; Thill et al., 2013; Tucker & Ellis, 1998). Neuroimaging and behavioral evidence has shown that merely observing manipulable objects can elicit activity in motor-related brain regions (Chao & Martin, 2000; Creem-Regehr & Lee, 2005), supporting the view that the sensorimotor system plays a key role even in passive visual contexts. Building on this view, more recent accounts suggest that the sensorimotor system does not operate in isolation but interacts flexibly with other cognitive systems, particularly semantic memory, during object interaction (Mizelle et al., 2013; Pulvermüller & Fadiga, 2010; Seidel et al., 2023). A number of studies have explored how visual attention is modulated by different types of object-tool interactions or intended actions, including passive viewing, grasping, and functional use (Ambrosini & Costantini, 2017; Borghi et al., 2012; Decroix & Kalénine, 2019; Juravle et al., 2015; Natraj et al., 2015; Tamaki et al., 2020). Concurrently, a growing body of literature has begun to highlight the importance of semantic knowledge in shaping the visual encoding of tools. These findings suggest that semantic expectations may influence attentional deployment and modulate the activation of action-related representations, particularly when objects are novel or conceptually ambiguous (Enge et al., 2023; Federico, Osiurak, Reynaud, et al., 2021; Federico, 2023; Federico, Osiurak, Brandimonte, et al., 2023; Federico & Brandimonte, 2019, 2020; Foerster, 2023; Foerster & Goslin, 2021; Lai et al., 2023; Pilacinski et al., 2021, 2021).

Visual object recognition is historically regarded as a hierarchical, multi-stage process. Early visual stages involve the extraction of basic features such as color, depth, and contour, which are subsequently grouped into meaningful shapes (Biederman, 1987). These perceptual representations are then matched with structural descriptions stored in long-term memory, where semantic attributes—such as object identity and familiarity—contribute to recognition. Familiarity, in particular, constitutes a core semantic dimension, reflecting the extent to which an object has been previously encountered or is known to the observer. It enables top-down modulation of visual processing, refining perception through learned expectations (Bar, 2021; Bar et al., 2006; Borghi et al., 2012; Connor et al., 2004; Decroix & Kalénine, 2019; Enge et al., 2023; Humphreys & Forde, 2001). However, most research in this domain has focused predominantly on highly familiar tools—objects with strong semantic associations and well-established functions. This raises an important question: Are the commonly reported effects of facilitated motor responses to tools genuinely due to action-related properties, or could they reflect a ceiling effect driven by overlearned semantic information?

To disentangle these possibilities, one may examine how unfamiliar tools—objects that afford action but lack prior semantic associations—are visually processed. Recent work by Federico, Osiurak, Brandimonte, et al. (2023) showed that repeated exposure to unfamiliar tools shifts gaze allocation from functional to manipulative components, suggesting a gradual integration of semantic knowledge with sensorimotor representations. Likewise, Juravle et al. (2015) observed an early attentional preference for functional features in novel objects. Yet, these studies focused exclusively on unfamiliar tools, precluding direct comparison with familiar ones. While other studies have attempted such comparisons (Belardinelli et al., 2015; Tamaki et al., 2020), their stimuli predominantly featured atypical familiar tools, rather than entirely novel ones. This over-representation potentially restricted the findings regarding how truly unfamiliar objects are explored in comparison to known ones. A more informative approach may, indeed, involve systematically contrasting gaze patterns toward truly novel versus well-known tools, thereby revealing how the presence—or absence—of familiarity modulates their visual exploration.

In line with the previously proposed “semantic-to-sensorimotor” cascade mechanism of tool-related visual processing (Federico & Brandimonte, 2019), we hypothesize that visual attention is initially directed toward semantic processing—namely, identifying the object and inferring its intended function—before transitioning to sensorimotor processing to assess how the object can be manipulated. This sequential allocation of attention is expected to be especially pronounced for unfamiliar tools, as semantic identification is likely to require greater cognitive effort, thereby shaping the overall pattern of visual exploration.

## Methods

### Participants

The study involved twenty-five right-handed participants (14 females; mean age = 23.6 years, SD = 3.35 years). All individuals provided written informed consent in strict accordance with the Declaration of Helsinki (1964). Participants were carefully screened through self-reports to ensure consistent right-handedness, normal or corrected-to-normal vision, and the absence of any history of neurological or psychiatric disorders. The sample size was determined based on a preliminary pilot study, employing an *a priori* power analysis to detect a medium effect size (η_p_^2^ =.25) within a repeated-measures ANOVA, with a statistical power of .80 and an alpha level of .05 (Faul et al., 2007, p. 200).

### Materials

The stimulus set consisted of twenty greyscale prehensile objects, equally divided into ten familiar and ten unfamiliar items. These stimuli were created using TinkerCAD and GrabCAD software. Images were displayed on a white background, measuring 1280x600 pixels, presented on a 1920x1080 grey screen. All images were rendered in grayscale. While the resolution varied, a constant length of 1080 pixels was maintained, with heights ranging from 140 to 511 pixels. The manipulative component was consistently positioned on the right and the functional component on the left across all stimuli. These components constitute the Areas of Interest (AOIs) subsequently considered in the analyses. All stimuli were centrally presented on a 27-inch monitor. Examples of both familiar and unfamiliar tools are depicted in *Figure 1*. An independent panel of ten university students (8 female, mean age 25.6 years, SD 3) was recruited to assess the appropriateness of the stimuli. These jurors evaluated each prehensile object for familiarity (5-point Likert scale) and right-hand graspability (binary “yes” or “no” response to the question: “Can you grasp this object with your right hand?”). This assessment aimed to confirm whether the tool’s structural characteristics afforded grasping and whether participants perceived the tool as manipulable. Familiarity responses below 2 (out of 5) for familiar prehensile objects were excluded, as they indicated insufficient familiarity (mean familiarity = 4.8, SD = 0.4). Conversely, responses above 2 for unfamiliar prehensile objects were excluded (mean familiarity = 0.7, SD = 0.3). For graspability, stimuli with scores below 50 per cent were excluded, as this threshold indicated they were perceived as non-graspable. Familiar prehensile objects achieved 100 per cent graspability (SD = 0), while unfamiliar prehensile objects demonstrated 85 per cent graspability (SD = 7%). Based on these criteria, no stimuli were ultimately excluded from the study, confirming their appropriate categorisation and perceived manipulability.

**Figure 1.**
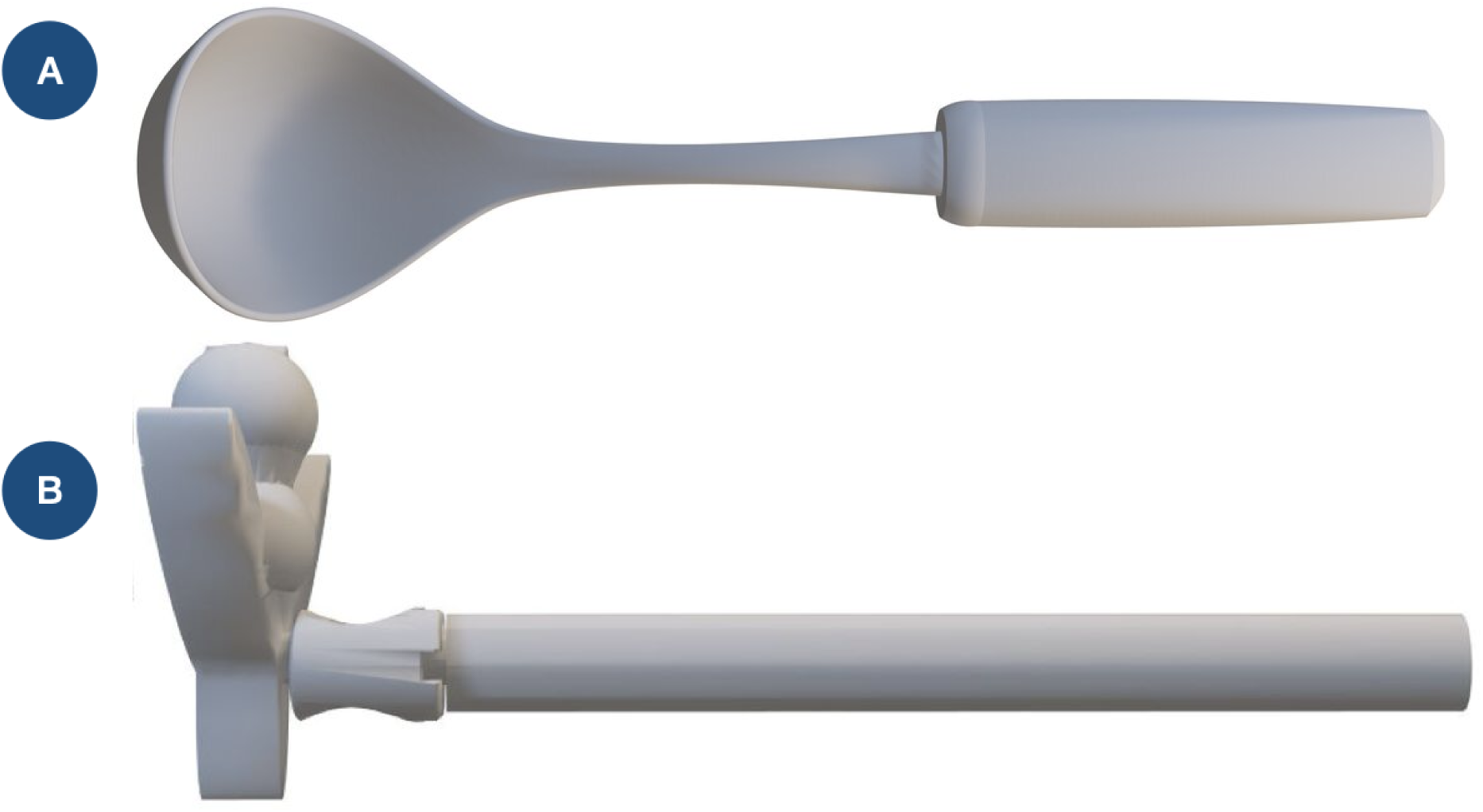
Stimuli. The study utilized twenty monochromatic images of graspable objects (ten familiar, ten unfamiliar) created in TinkerCAD and GrabCAD. All images displayed the manipulation area on the right and the functional area on the left. Example stimuli included (A) a familiar object and (B) an unfamiliar object.

### Procedure

The study was conducted at the Laboratory of Experimental Psychology at Suor Orsola Benincasa University in Naples, Italy (https://www.cogsci.it). Upon arrival, participants provided written informed consent and were then seated in front of the eye-tracking system. To minimise head movement and ensure high-precision eye movement recording, participants were instructed to rest their chins and foreheads on a chin rest. They were positioned approximately 64 cm from the monitor, with their right hand resting visibly motionless on the desk. Before the experimental session, participants received detailed task instructions and completed a standard nine-point eye-tracking calibration procedure (Gibaldi et al., 2017). Following successful calibration, they were instructed to “observe what appears on the screen as naturally as possible,” after which the experiment commenced. The study employed a 2 (familiar vs. unfamiliar tools) x 2 (Function AOI vs. Manipulation AOI) within-subjects design. Each trial adhered to a fixed visual sequence: first, a black fixation point appeared centrally on a grey screen for 500 ms; then, the stimulus was presented for 1500 ms; finally, a grey blank screen was displayed for 3000 ms. The total duration of each trial was 5000 ms. Each tool was presented four times in random order (*i.e.,* four repetitions), resulting in a total of 80 trials per participant. The total eye-tracking session duration of approximately ten minutes per participant. The experimental flow is visually represented in *Figure 2*. After the experiment, participants completed a questionnaire evaluating the familiarity, graspability, and usability of the objects. For the usability judgement, they responded “yes” or “no” to the question: “Would you use this object with your right hand to act?” If they selected “no,” they were prompted to provide an explanation. Finally, participants were debriefed regarding the study’s purpose.

**Figure 2.**
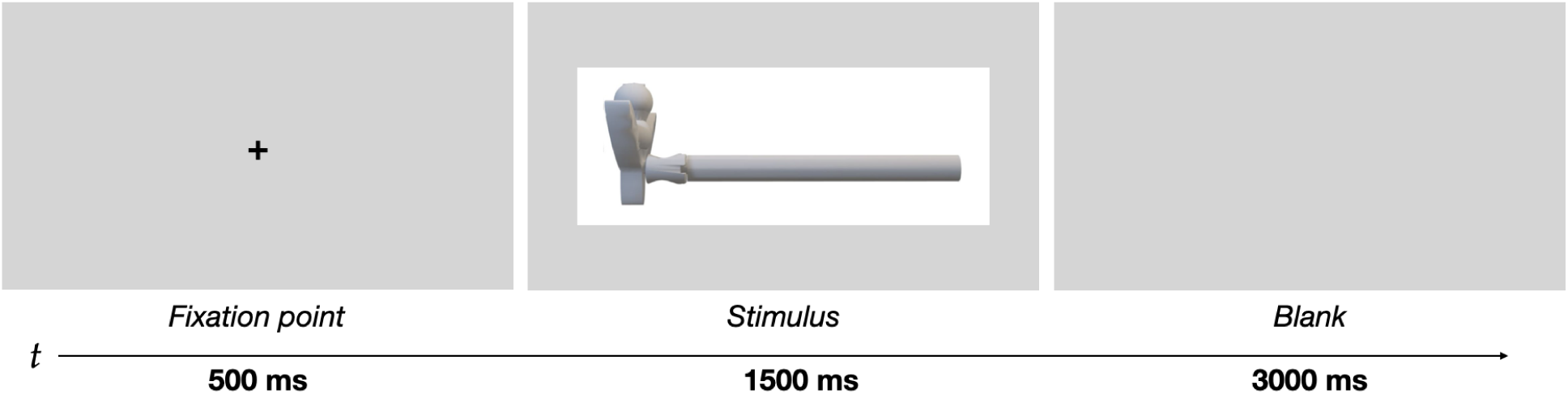
Experimental flow. Participants were presented with a fixation point for 500 ms, followed by a stimulus displayed for 1500 ms. Subsequently, a blank screen was shown to allow for eye relaxation. Eye movements were recorded throughout the entire experimental procedure.

### Apparatus

Gaze-behaviour data were acquired using a Tobii EyeX eye-tracking system, operating at a sampling rate of 60 Hz, interfaced via Myex software (Jones, 2018). All experimental paradigms were conceived and programmed using the MATLAB R2024b.

### Eye-tracking Data Preprocessing

The proportion of gaze samples (*x*,*y* coordinates) falling within each AOI was analyzed to understand participants’ visual exploration. To avoid misclassifying small eye movements, the AOIs were defined broadly, extending 100 pixels horizontally from the image’s center to the edges. In the vertical plane, AOI height was determined by the maximum height of the stimulus, ensuring that the full extent of the object was encompassed within the defined areas. Examples of these AOIs are shown in *Figure 3*. Preliminary inspection of the gaze data revealed a systematic leftward shift in horizontal fixation positions across participants, which may be attributable to a measurement error in the eye-tracking system. This was statistically confirmed by a one-sample *t-test* conducted on the average horizontal fixation coordinates recorded between 200 and 500 ms from trial onset—thus excluding the initial refixation phase. Results showed that the mean gaze position was significantly left of the screen center (*M* = –136.15 px, *t*(24) = –9.6, p < .001). To correct this bias, the average gaze position (*x*, *y* coordinates) between 200 ms and 500 ms of viewing was computed across all participants and trials. This average offset was then subtracted from the gaze coordinates of all subsequent time points in each trial, effectively re-centring the data. For clarity, while the nominal screen centre was at (960, 540) pixels, after this correction, the new origin was defined such that the corrected mean gaze position aligned with (0, 0).

**Figure 3.**
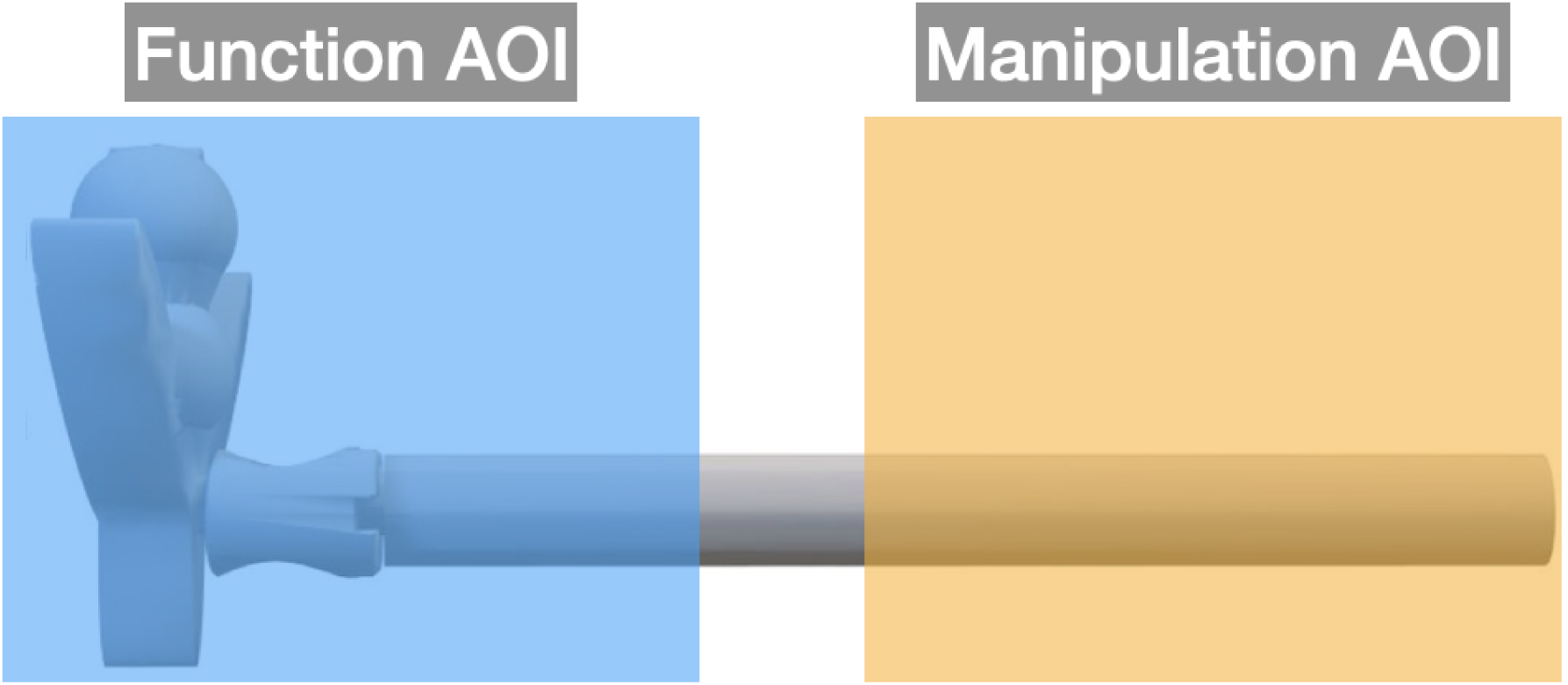
Areas of Interest (*AOIs*). The Functional AOI (blue rectangle) and the Manipulation AOI (yellow rectangle). To avoid misclassifying minor saccades near the fixation point, each AOI extended 100 pixels horizontally from the image’s center to its edges.

### Data Analysis

To characterize the temporal structure of gaze allocation as a function of familiarity, a two-step analytical procedure was implemented (Federico, Osiurak, Brandimonte, et al., 2023). Initially, a repeated-measures analysis of variance (ANOVA) was conducted with Areas of Interest (*AOI*: functional, manipulation) and *Familiarity* (familiar, unfamiliar) as within-subject factors. The dependent variable was the proportion of gaze samples (*x*,*y* coordinates) falling within each AOI, aggregated across conditions (hereafter, gaze samples). This measure captures the overall spatial distribution of visual attention, thus reflecting the full temporal profile of viewing behavior. Bonferroni-adjusted pairwise comparisons were used to test specific contrasts of interest, controlling for family-wise error rate (Benjamini & Hochberg, 1995).

To further model the temporal dynamics of gaze behavior across the entire 1500-millisecond stimulus presentation window, a growth curve analysis (GCA) was implemented using linear mixed-effects models parameterized with orthogonal polynomials (Mirman, 2017). For each *AOI*, time was modeled as a continuous predictor and decomposed into three mutually orthogonal polynomial terms: linear (*ot1*), quadratic (*ot2*), and cubic (*ot3*), computed via the Gram-Schmidt orthonormalization process. The first-order polynomial (*ot1*) represents the linear trend over time, capturing monotonic increases or decreases in gaze allocation to a given AOI. The second-order polynomial (*ot2*) accounts for quadratic, symmetric nonlinear trends, such as an increase followed by a decrease, reflecting a single inflection point. The third-order polynomial (*ot3*) models cubic curvature, allowing for more complex non-monotonic dynamics with two inflection points—such as patterns where gaze allocation increases, then decreases, and increases again. The model’s structure was defined as:

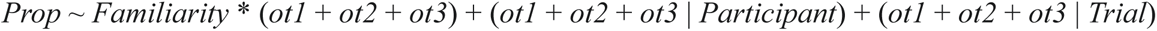

where *Prop* indicates the proportion of gaze samples within the *AOI* at each time point. The model included fixed effects for *Familiarity*, linear and higher-order temporal terms, and their interactions, as well as random intercepts and slopes for all time components at both the participant and trial levels. To statistically evaluate these temporal bends and inflection points, an ANOVA for mixed-effects models was performed using Satterthwaite’s method for degrees of freedom approximation (Kuznetsova et al., 2017). A significance threshold of α = 0.05 was adopted for all tests, and Bonferroni correction was applied to adjust for multiple pairwise comparisons. All statistical analyses were conducted in R (v. 4) on Apple macOS Sonoma 14.6.

## Results

Descriptive statistics are reported in *Table 1*.

**Table 1.**
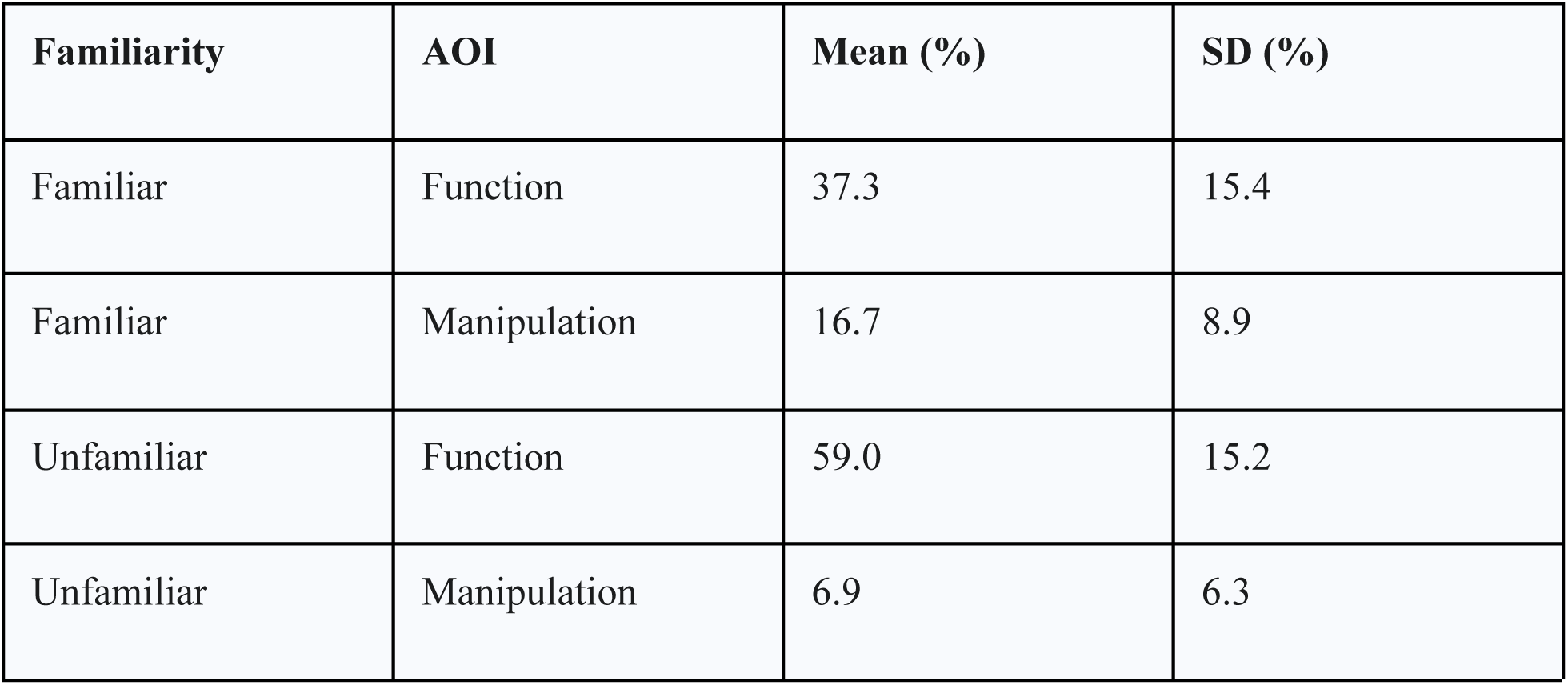
Mean and standard deviation (SD) of proportion of gaze samples during the overall visual exploration (1500 ms).

The repeated-measures ANOVA revealed a significant main effect of *AOI* (F(1, 360) = 1203.2, p < .001, η_p_² = .77). A main effect of *Familiarity* also emerged (F(1, 360) = 32.81, p < .001, η_p_² = .08). Finally, a significant interaction between *AOI* and *Familiarity* was observed (F(1, 360) = 225.21, p < .001, η_p_² = .38). Bonferroni-corrected post hoc analyses confirmed that, across both familiar and unfamiliar objects, the proportion of gaze samples on the functional *AOI* was significantly greater than that on the manipulation *AOI*. Also, participants spent a higher proportion of gaze samples to the functional *AOI* when the tools were unfamiliar, whereas the manipulation *AOI* attracted greater attention when the tools were familiar. The results of the repeated-measures ANOVA are summarized in *Figure 4A*.

**Figure 4.**
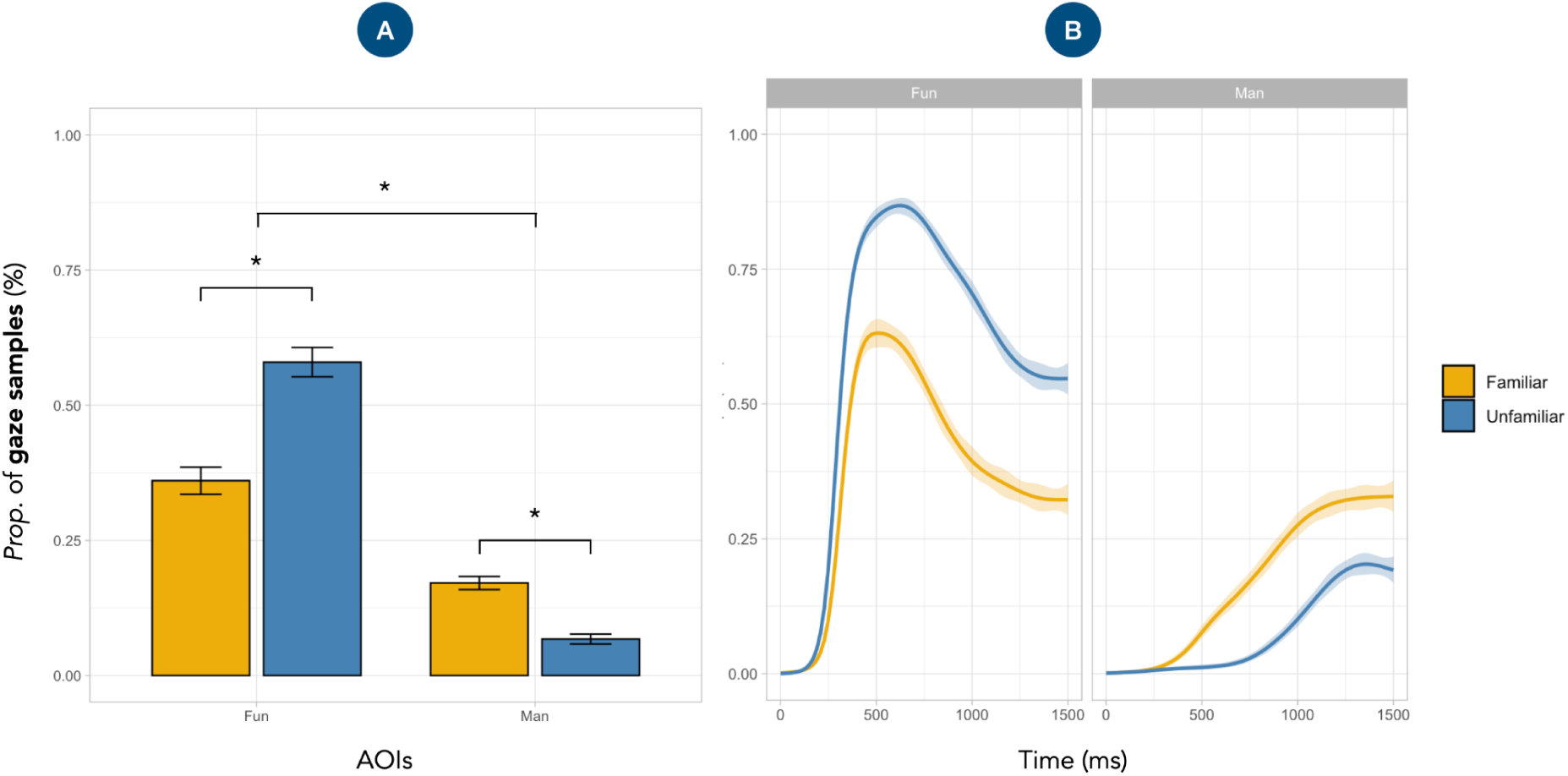
Results. (A) Results of the repeated-measures ANOVA showing the proportion of gaze samples directed to functional (Fun) and manipulative (Man) areas of interest (AOIs) as a function of tool familiarity. Significant main effects of *AOI* and *Familiarity* were observed, as well as a significant *AOI* × *Familiarity* interaction. Overall, participants fixated more on functional than manipulative components, with increased attention to functional regions for unfamiliar tools and to manipulative regions for familiar tools. Error bars indicate standard errors. * = *P* < .001. (B) Time-course of gaze allocation over a 1500-ms viewing window, modeled via Growth Curve Analysis, for each AOI and condition. Shaded areas represent 95% confidence intervals.

The GCA provided a temporally resolved account of how tool familiarity modulates gaze behavior over time. For the functional *AOI*, there was a significant main effect of *Familiarity*, *F*(1, 44,502) = 3226.50, *p* < .001, and a significant effect of the cubic time term *ot3*, *F*(1, 32) = 79.95, *p* < .001. The interaction between *Familiarity* and *ot3* was also significant, *F*(1, 40,660) = 7.29, *p* = .007. Additional significant effects were found for *ot1*, *F*(1, 32) = 75.26, *p* < .001, and *ot2*, *F*(1, 27) = 230.76, *p* < .001, as well as for the interactions *Familiarity* × *ot1*, *F*(1, 43,772) = 290.67, *p* < .001, and *Familiarity* × *ot2*, *F*(1, 41,328) = 347.19, *p* < .001. For the manipulation *AOI*, there was a significant main effect of *Familiarity*, *F*(1, 43,286) = 1161.70, *p* < .001. Significant effects were also observed for *ot1*, *F*(1, 33) = 87.27, *p* < .001, and *ot3*, *F*(1, 34) = 12.61, *p* = .001, while the effect of ot2 did not reach significance, *F*(1, 35) = 2.92, *p* = .096. All interaction terms were significant: Familiarity × *ot1*, *F*(1, 44,080) = 339.82, *p* < .001; Familiarity × ot2, *F*(1, 41,532) = 105.96, *p* < .001; and Familiarity × *ot3*, *F*(1, 34,459) = 21.10, *p* < .001. Full GCA results are presented in *Figure 4B*.

Taken together, the results reveal that tool familiarity shapes both the overall distribution and the temporal dynamics of visual attention. The repeated-measures ANOVA showed that participants allocated more gaze samples to the functional *AOI* than to the manipulative *AOI*, and that this allocation varied as a function of familiarity. Specifically, the significant *AOI* × *Familiarity* interaction indicated that attention to the functional region was greater for unfamiliar tools, while gaze samples to the manipulative region increased for familiar tools. The GCA complemented this analysis by providing a temporally resolved profile of gaze behavior. Significant effects for the linear (*ot1*), quadratic (*ot2*), and cubic (*ot3*) time terms, along with their interactions with *Familiarity*, indicated that a third-order polynomial best captured the non-linear trajectory of gaze allocation over time. Attention initially prioritized functional features, but progressively shifted toward manipulative components—particularly for familiar tools—suggesting a sequential transition from semantic analysis to action-related processing. The cubic *Familiarity* × *ot3* interaction reflected inflection points in gaze trajectories, potentially indexing attentional reinvestment during prolonged observation (Busjahn & Tamm, 2021). These reallocations may result from increased semantic demands or stimulus-driven salience effects.

## Discussion

This eye-tracking study examined the visual exploration patterns of participants engaged in a free-viewing task, aiming to investigate how familiarity influences the allocation of visual attention to functional and manipulation AOIs of both familiar and unfamiliar tools. Results revealed distinct patterns of gaze allocation, which were modulated by the familiarity of the tools. During the initial phase of visual exploration, participants were focused on the functional component of tools, irrespective of their familiarity. This initial attentional bias suggests that observers may prioritise the extraction of semantic information related to the object’s identity and purpose. For unfamiliar objects, this effect was particularly prominent, with participants dedicating almost no time to the manipulation AOI in the first 500ms of visual exploration.

The gaze patterns we found align with recent evidence suggesting the critical role of semantic processing in tool visual exploration (Enge et al., 2023; Federico, Osiurak, Reynaud, et al., 2021; Federico, Osiurak, Brandimonte, et al., 2023; Federico & Brandimonte, 2019; Foerster & Goslin, 2021; Juravle et al., 2015). In this epistemological perimeter, it is only after gaining a preliminary understanding of the object’s identity and function that attention shifts towards its manipulation components, presumably in preparation for potential action (Bayani et al., 2021; Enge et al., 2023; Foerster & Goslin, 2021). Recent studies similarly observed an initial visual exploration of tool–related functional components, with a shift towards manipulative parts only as familiarity increased (Federico et al., 2022; Juravle et al., 2015). Globally, this study’s results support the recently proposed “semantic-to-sensorimotor” cascade mechanism (Federico, Osiurak, & Brandimonte, 2021; Federico, Osiurak, Brandimonte, et al., 2023; Federico, Osiurak, Ciccarelli, et al., 2023; Federico, Osiurak, Reynaud, et al., 2021; Federico & Brandimonte, 2019, 2019, 2020). Additionally, the sustained attention to functional aspects of unfamiliar tools aligns with object recognition studies showing longer fixations indicate difficulty in semantic extraction (Just & Carpenter, 1976). Taken together, the results support the “action-reappraisal” hypothesis, which suggests that the ease of integrating sensorimotor and semantic knowledge affects action planning ability (Federico & Brandimonte, 2019).

A key aspect of the current study is the consistent right-hand orientation of manipulation components for all tools—including unfamiliar ones—and the exclusive participation of right-handed individuals. This design represents a stringent test for hypotheses predicting an immediate prioritization of affordance-driven manipulative features (Thill et al., 2013). Specifically, if the visual system preferentially processed action-related information, the positioning should have elicited rapid attention to the manipulation/graspable area, priming motor responses (Tucker & Ellis, 1998). However, contrary to this prediction, the growth curve analysis revealed that initial attention to manipulation areas was almost non-existent for unfamiliar tools in the first 500ms of viewing, increasing slowly and remaining relatively low as compared to familiar tools. This delayed attention towards the manipulation features of unfamiliar tools suggests that semantic understanding might be prioritized initially. This, in turn, supports the idea that the visual system may prioritize identifying objects and understanding their purpose before focusing on immediate motor affordances (Enge et al., 2023).

An alternative explanation for the findings of this study involves the role of perceptual salience in driving early attentional capture (Nuthmann et al., 2020; Spotorno et al., 2013). Highly salient visual features tend to attract early fixations. Therefore, it is plausible that the functional components of the tools—particularly in the case of unfamiliar objects—appeared more perceptually salient due to their atypical or novel visual characteristics. While such bottom-up salience may indeed contribute to the initial capture of attention, it is inherently a transient effect and thus cannot account for the sustained engagement observed in the functional AOIs (Spotorno & Tatler, 2017). In the present data, prolonged attentional allocation to these regions—especially for unfamiliar tools—is more plausibly attributed to top-down, semantically driven processing (Federico, Osiurak, Ciccarelli, et al., 2023; Federico & Brandimonte, 2020). Nevertheless, because perceptual salience was neither quantified nor experimentally controlled in the current design, both bottom-up and top-down interpretations remain viable. In this context, the GCA becomes particularly informative: the presence of two inflection points in the trajectory of the cubic temporal term may reflect a shift from early salience-based orienting to later semantic elaboration. This is consistent with theoretical accounts that describe object perception as a cascade of early visual filtering followed by conceptual integration (Enge et al., 2023; Federico, 2023). Future studies employing complementary paradigms—such as forced-choice tasks or fixed-gaze designs—may help clarify the relative contributions of these mechanisms. Additionally, combining eye-tracking with physiological measures like electromyography and electroencephalography could enable the co-registration of motor preparation and cortical activity during the visual exploration of familiar versus unfamiliar tools.

## Conclusion

This eye-tracking study investigated how familiarity with tools modulates their visual exploration during free viewing. Observers consistently prioritized functional components, particularly when tools were unfamiliar, indicating that initial visual attention might be guided by semantic demands. In contrast, gaze samples on manipulation-related features emerged later and were more pronounced for familiar tools, likely reflecting reduced semantic load and more rapid access to action-related information. These findings support emerging accounts of action understanding that emphasize a semantic-to-sensorimotor cascade mechanism in the visual encoding of tool-related information (Enge et al., 2023; Federico & Brandimonte, 2020).

## Author contributions

G.F. conceived the study. L.V. and M.C. implemented the eye-tracking paradigm and conducted the experiment. L.V. performed the data analysis. C.R.I. provided support in data analysis. G.F. wrote the manuscript with input from all the authors. All the authors approved the manuscript.

## Acknowledgements

The authors thank Professor Antimo Buonocore (Suor Orsola Benincasa University, Naples, Italy) for his support in developing the eye-tracking paradigm, for his comments, and for his assistance with the analyses.

## Data analysis

The data supporting the present study’s findings are available at https://osf.io/8ptqj/.

